# Population diversity jeopardizes the efficacy of antibiotic cycling

**DOI:** 10.1101/082107

**Authors:** Yunxin J. Jiao, Michael Baym, Adrian Veres, Roy Kishony

## Abstract

Treatment strategies that anticipate and respond to the evolution of pathogens are promising tools for combating the global rise of antibiotic resistance^1–3^. Mutations conferring resistance to one drug can confer positive or negative cross-resistance to other drugs^4^. The sequential use of drugs exhibiting negative cross-resistance has been proposed to prevent or slow down the evolution of resistance^5–8^, although factors affecting its efficacy have not been investigated. Here we show that population diversity can disrupt the efficacy of negative cross-resistance-based therapies. By testing 3317 resistant *Staphylococcus aureus* mutants against multiple antibiotics, we show that first-step mutants exhibit diverse cross-resistance profiles: even when the majority of mutants show negative cross-resistance, rare positive cross-resistant mutants can appear. Using a drug pair showing reciprocal negative cross-resistance, we found that selection for resistance to the first drug in small populations can decrease resistance to the second drug, but identical selection conditions in large populations can increases resistance to the second drug through the appearance of rare positive cross-resistant mutants. We further find that, even with small populations and strong bottlenecks, resistance to both drugs can increase through sequential steps of negative cross-resistance cycling. Thus, low diversity is necessary but not sufficient for effective cycling therapies. While evolutionary interventions are promising tools for controlling antibiotic resistance, they can be sensitive to population diversity and the accessibility of evolutionary paths, and so must be carefully designed to avoid harmful outcomes.

## Main Text

The use of antibiotic combinations can impede the evolution of resistance by imposing evolutionary tradeoffs^3^. Recent attention to these phenomena has been driven by the promise of developing more effective strategies to prevent treatment failure due to de novo mutations in long-term infections, by exploiting the evolutionary interactions between drugs to impede the development of resistance^5–7,9–13^. In general, mutations providing resistance to one drug can also either increase or decrease resistance to other drugs. While increased resistance to other drugs (positive cross-resistance) is common^5,6^, for some drug pairs, resistance to one drug decreases resistance to another drug (negative cross-resistance, also called collateral sensitivity^7^). Systematic studies have revealed specific drug pairs with negative cross-resistance, including cases of reciprocal negative interactions, whereby resistance to any of the two drugs confers sensitivity to the other^4–6,14^. Simultaneous use of drug pairs with negative cross resistance has been explored as a means to slow down the evolution of resistance to toxins ranging from antibiotics to pesticides^5–7,15–18^. Alternatively, drugs with collateral sensitivity can be used sequentially to slow evolution of resistance, thus avoiding toxicity or incompatibility issues that complicate antibiotic mixtures in clinical settings^5,7^.

It remains unknown whether the effectiveness of alternating negative cross-resistance treatment is robust to the large population size often found in clinical infections, and to what extent longer-term evolution can be reliably predicted by the cross-resistance profiles of first-step mutants. Previous measurements on negative cross-resistance have typically assayed a small number of first-step mutants, and evolution experiments were conducted with small population sizes^5–7,9,19,20^. Because the diversity of cross-resistance across random mutants has not been systematically measured, it is unknown whether negative cross-resistance seen in typical mutants is representative, and whether negative cross-resistant treatment would be effective for larger populations with larger potential for rare mutants.

We focus here on *Staphylococcus aureus*, a common human pathogen in which *de novo* antibiotic resistance is of clinical importance^21–25^. We developed a high-throughput assay to profile the cross-resistances of 3317 resistant mutants to six antibiotics with diverse mechanisms of action: oxacillin (OXA), novobiocin (NOV), ciprofloxacin (CPR), gentamicin (GEN), amikacin (AMK), and doxycycline (DOX) (Fig. 1a; **Supplementary Table 1**).

**Fig. 1.**
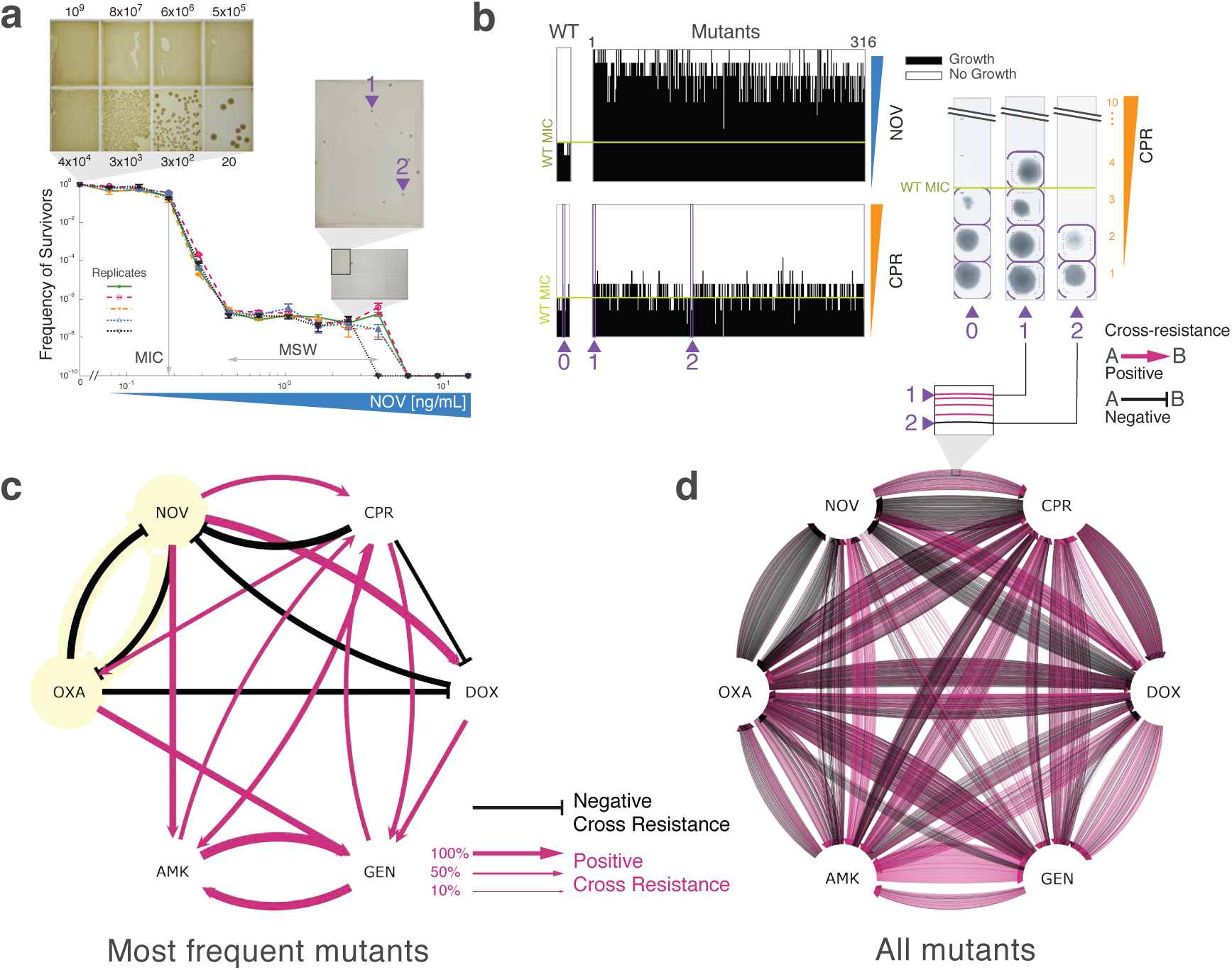
High-throughput measurement of cross-resistance in 3,317 *de novo* mutants reveal considerable diversity in both the magnitude and sign of cross-resistance. a) The mutant selection window for NOV was measured by counting the frequency of survivors at 10 drug concentrations. Each line indicates an independent biological replicate. Inserts show examples from the automated imaging pipeline. b) Measured resistance to NOV and cross-resistance to CPR of mutants selected on NOV, showing a range of MICs for both drugs. Examples of the automated image pipeline are shown for two isolates (1 and 2), chosen as indicated from the inset in (a). c) The network of most frequently seen cross-resistance interactions between every drug pair in our library. The majority of pairs show positive cross-resistance. Only the most common phenotype out of three possible phenotypes (positive, negative, and neutral cross-resistance) representing a plurality (>33%) of all mutants are shown. The one reciprocal negative cross-resistant pair, OXA and NOV, is highlighted. d) The full network of cross-resistance for all observed mutants, showing heterogeneous interactions for all pairs. Here every mutant is represented as a single edge originating from the drug it was selected in.

The frequency of surviving colonies as a function of antibiotic concentration follows a characteristic shape: survival falls off when drug concentration exceeds the minimum inhibitory concentration (MIC), and then plateaus around 10^–6^-10^-8^ until it drops below detection at drug concentrations high enough to kill even the most resistant mutant (Mutant Prevention Concentration, MPC)^26^. The plateau region between the MIC and MPC is termed the mutant selection window (MSW), and constitutes the range of antibiotic concentrations where selection for resistance occurs. For each antibiotic, we collected roughly 500 resistant mutants from plates in the MSW range (Fig. 1a; **Supplementary Fig. 1; Methods**). We measured the level of resistance of each of these mutants to each of the 6 antibiotics by growing the library on a series of agar plates at increasing antibiotic concentrations (Fig. 1b; **Methods**). Positive cross-resistance was detected when a mutant survived at higher drug concentrations than the wild-type, while negative cross-resistance was indicated by the wild-type growing at a higher drug concentration than the mutant. (Fig. 1b). Using a robotic pipeline, we systematically measured cross-resistance interactions in nearly 20,000 mutant-drug combinations.

To provide a high-level picture of cross-resistance interactions and identify negative-cross resistance pairs, we constructed a network of the modal cross-resistance between drug pairs. Out of the 30 possible ordered drug pairs, our analysis revealed 6 negative and 12 positive cross-resistance interactions. The negative cross-resistance interactions included the reciprocally cross-resistant pair, OXA and NOV, and the three-drug negative cross-resistant cycle, OXA->DOX->NOV->OXA. In contrast to the negative cross-resistance observed between aminoglycosides and other drugs in *E. coli*^5,6^, we found predominantly positive cross-resistance interactions between aminoglycosides and other antibiotics (Fig. 1c). These observations highlight the species-specificity of cross-resistance interactions, and indicate that cross-resistance-based treatment strategies must be tailored for specific organisms.

### Cross-resistance profiles are highly heterogeneous

Analysis of the distribution of cross-resistance reveals that, for all drug pairs, there is considerable variation in magnitude and sign of cross-resistance among the collection of resistant mutants. Even for drug pairs where mutants primarily display cross-resistance of a specific sign, there are rare mutants with phenotypes opposite to the general trend (Fig. 1d; **Supplementary Fig. 2a**). The MICs of the two aminoglycosides, AMK and GEN, were positively correlated, however we observed no strong MIC correlations between antibiotic pairs from different classes (**Supplementary Fig. 3**). Going beyond the typical measurements of small numbers of resistant mutants revealed hitherto unseen diversity in cross-resistance profiles (**Supplementary Fig. 3**).

To investigate the impact of diversity in cross-resistance, we focused on the reciprocal negative cross-resistance pair OXA and NOV (Fig. 1b). While for the majority of OXA and NOV resistant mutants, negative cross-resistance to the other drug was observed, rare, positive cross-resistant mutants also appear (Fig. 2). Of the OXA-resistant mutants, 74% have negative cross-resistance to NOV, 21% do not affect NOV resistance, while 5% have positive cross-resistance. Of the NOV-resistant mutants, 62% exhibit negative cross-resistance to OXA, 33.5% do not affect OXA resistance and 4.5% display positive cross-resistance. These observations suggest that while small-scale studies may identify OXA and NOV as a reciprocal negative cross-resistance pair, they would miss the underlying fraction of rare mutants with positive cross-resistance, which could in turn affect the evolution of multi-drug resistance.

**Fig. 2.**
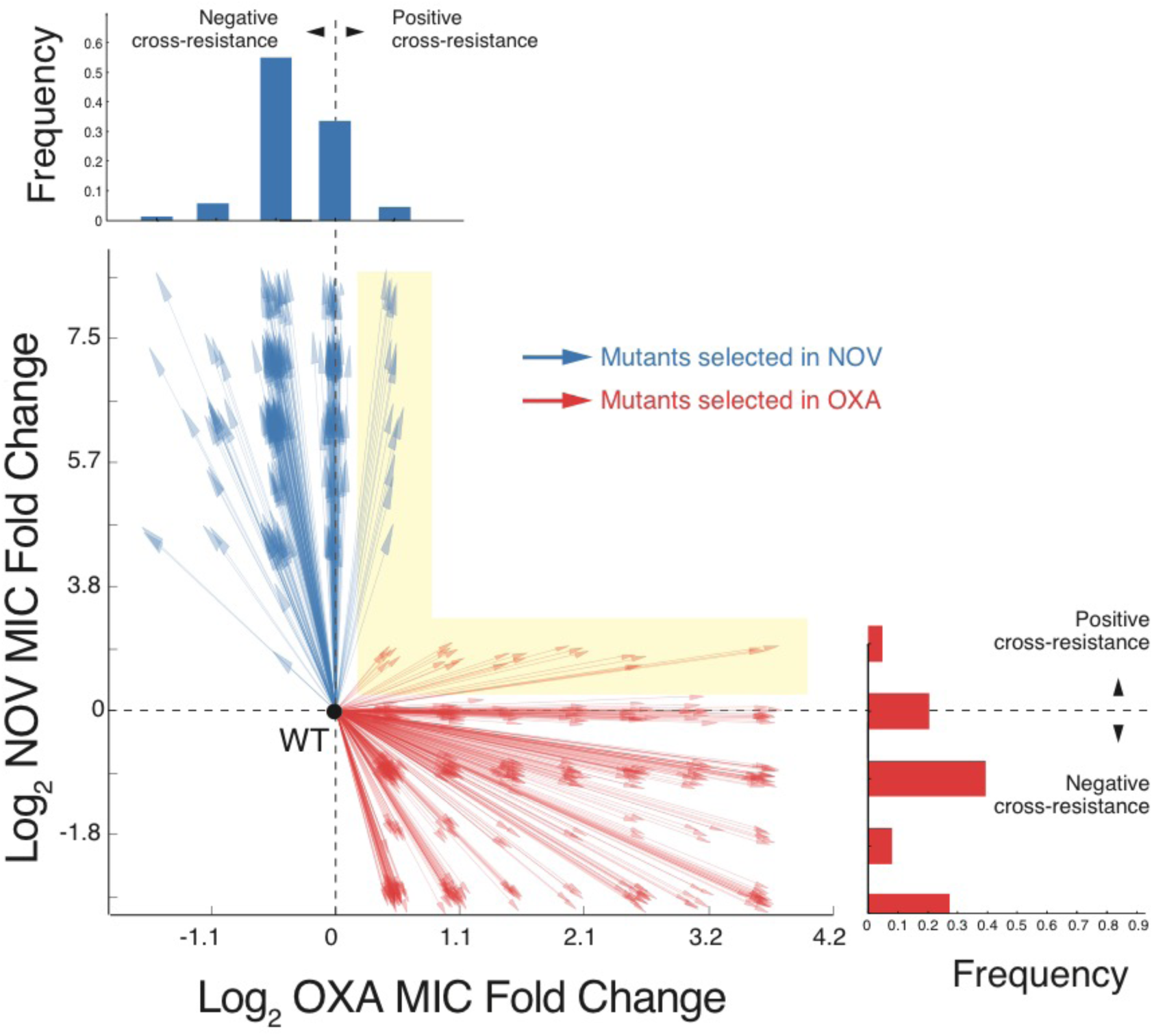
Reciprocal negative cross-resistance between OXA and NOV is disrupted by rare positive cross-resistant mutants. Vectors indicate trajectories of one-step mutants on the 2D plane defined by OXA and NOV resistance. Histograms display the distribution of cross-resistance phenotypes. Red, mutants selected in OXA; blue, mutants selected in NOV. Rare, positive cross-resistant mutants are highlighted.

### Rare mutants appearing in large populations reverse the outcome of cycling

We next asked whether rare cross-resistant phenotypes can impact the effectiveness of negative cross-resistance treatments. As rare positive cross-resistant mutants are more likely to appear in larger populations, we hypothesized that the outcome of sequential treatment with a reciprocal negative cross-resistance drug pair would be affected by population size. We first considered treatment with OXA followed by NOV, followed by the reciprocal. Based on the measured frequencies of resistance (**Supplementary Fig. 1**), we selected for resistant mutants at the same concentration from either a small (10^6^ cells) inoculum, in which approximately one mutant is expected to survive, and a large (10^9^ cells) inoculum, in which approximately a thousand mutants are expected to survive. After selection for resistance to the first drug, we expanded the two populations to the same size, and measured the frequency of survivors as a function of the concentration of the second drug, as compared to the ancestral wildtype (**Methods**, Fig. 3a).

**Fig. 3.**
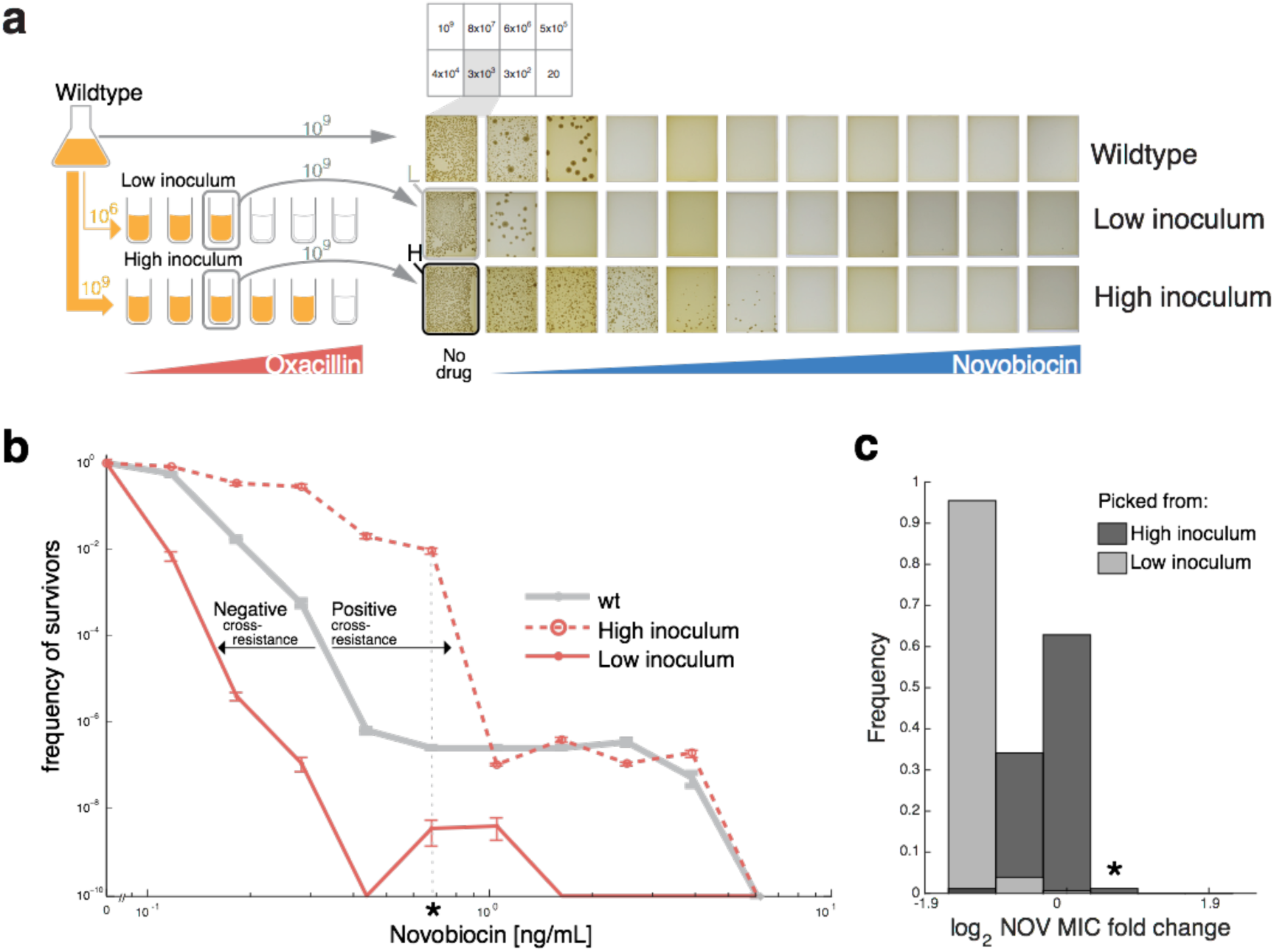
Negative cross-resistance cycling selects for positive cross-resistance in large populations due to appearance of rare positive cross-resistant mutants. a) A wild type population was split into a large inoculum (10^9^) or small inoculum (10^6^) and selected in identical concentrations of OXA. The two populations, along with the ancestor, were then grown to the 1e9 cells/mL before plating on NOV. b) The low diversity population (dashed red line) exhibits negative cross-resistance. However, the high diversity population (solid red line), which differs from the low diversity population only in the starting population size, exhibits positive cross resistance. c) NOV-resistance profiles of colonies were picked from the high and low inoculum populations after selection in oxacillin. The low inoculum population (L) show primarily negative cross-resistance to NOV, while the high inoculum population (H) displays a much more diverse cross-resistance profile, including rare mutants that have positive cross-resistance to NOV.

While the low-inoculum OXA-evolved population exhibited marked negative cross-resistance to NOV, the high-inoculum population displayed not only a reduction but a complete inversion of this negative cross-resistance (Fig. 3b). The MIC of the low-inoculum population is reduced in comparison to the WT, demonstrating the efficacy of negative cross-resistance in using one drug in order to reduce resistance to another (fewer than one in 10^5^ survivors at 0.15 ng/mL NOV compared to WT 0.36 ng/mL). Beyond a reduction in MIC, this low-inoculum population also has lower potential to evolve as evident by its significantly reduced MPC (1.1 ng/mL NOV for the OXA-evolved population vs 4.0 ng/mL for the WT). In striking contrast, the population that was selected on the same exact drug concentration of OXA but with large inoculum size showed an increase rather than decrease in resistance (fewer than one in 10^5^ survivors at 0.86 ng/mL). Further, unlike the low-inoculum population, the high-inoculum population has the same MPC as the wild type, showing no reduction in evolutionary potential. The reciprocal case of NOV followed by OXA resulted in similar, albeit less drastic, inversion of negative cross-resistance due to inoculum size (**Supplementary Fig. 4a**).

Phenotyping mutants recovered after liquid selection, we found that the low inoculum population shows homogenous negative-cross resistance phenotypes while the high inoculum population is more diverse and contains rare positive cross-resistance mutants. Isolation of colonies from the low diversity population after OXA selection but before NOV selection shows a relatively homogenous population with negative cross-resistance to NOV (Fig. 3c, **L**). In contrast, colonies from the high inoculum population before NOV selection shows a wider spread of cross-resistance phenotypes (Fig. 3c, **H**). Only 1% of this high inoculum population showed positive cross-resistance with 2.4-fold increase in MIC over the wild type (Fig. 3c, **H**), in agreement with the sharpest drop-off in survival occurring at 1 in 10^2^ cells, at roughly 2.4 fold increase over the effective wild type MIC (Fig. 3b). Together, these results show that rare positive-cross resistant mutants can disrupt, and even invert the effect of negative cross-resistance cycling.

### Distinct mutation profiles mediate cross-resistance phenotypes

To understand the genetic mechanisms behind resistance and cross-resistance, we whole-genome sequenced 277 mutants that had been selected for resistance to OXA or NOV (see methods). NOV mutations were clustered around known resistance hotspots in *gyrB*^27^, while OXA mutations targeted general stress response pathways through binding partners of sigma factor *sigB*^28^ (p = 1.7⋅10^-4^, fisher exact test, **Supplementary table 2**). In the OXA mutants, 30 mutations occurred in *gdpP*, including 14 nonsense mutations (**Supplementary table 3, supplementary Fig. S5**). *gdpP* encodes a recently characterized putative c-di-AMP phosphodiesterase, and knockouts of *gdpP* have been shown to increase beta-lactam resistance^29–31^. We identified five mutations in relA, a gene in which point mutations have been linked to Linezolid resistance by switching on the stringent response in clinical isolates^25^. We also recovered mutations in fmt and penicillin-binding protein pbp1, both of which are involved in cell wall synthesis^21,32^ (**supplementary table 3**). The observation that OXA mutations target different pathways while NOV mutations target a single gene may explain why OXA resistance occur roughly 10 times more frequently than NOV resistance.

For each mutant, we then tested for association with different cross-resistance phenotypes (**supplementary Fig. S5**). In NOV mutants, *gyrB* R144I was associated with neutral cross-resistance (enrichment, p = 1.09·10^-4^, 2-tailed fisher exact test, Bonferroni adjusted) and negative cross-resistance (depletion, p=0.01). In OXA mutants, associations were found between *gdpP* E550* and negative cross-resistance (p = 3·10^-2^); rpoB G827D with both negative cross-resistance (depletion, p = 1.93·10^-4^) and neutral cross-resistance (enrichment, p = 2.69·10^-5^); and *rpoC* V78I with both negative cross-resistance (depletion, p = 2.61·10^-3^) and neutral cross-resistance (enrichment, p = 5.83·10^-4^). Due to the small number of mutants with positive cross-resistance, after correcting for multiple hypothesis testing, no mutations were significantly associated with positive cross-resistance. However, we recovered nonsynonymous mutations in *rsbW*, the negative regulator of the stress response sigma factor *sigB*^33^, only in OXA mutants with positive (n=2) or neutral cross-resistance (n=1) to NOV, in concurrence with the role of *rsbW* mutations in multiple drug resistance^7^. Taken together, we find that *in vitro* evolution recovers clinically important mutations, and that mutation profiles differ between cross-resistance phenotypes.

### Negative cross resistance is not preserved through sequential antibiotic selection

Our results suggest that negative cross-resistance cycling may be more successful if the population size was kept small at each step. To study whether maintenance of small population size is sufficient for success of negative cross-resistance cycling, we sequentially treated populations with OXA and NOV, transferring 10^6^ or 10^7^ cells at each step for OXA and NOV selection, respectively, to limit diversity based on the frequency of mutation (**Fig. S1**). We found that the first step consistently produces negative cross-resistance. However, as mutations accumulate in subsequent steps, negative cross-resistance frequently disappeared (Fig. 4). Thus, small population size appears necessary but not sufficient for successful negative cross-resistance cycling.

**Fig. 4.**
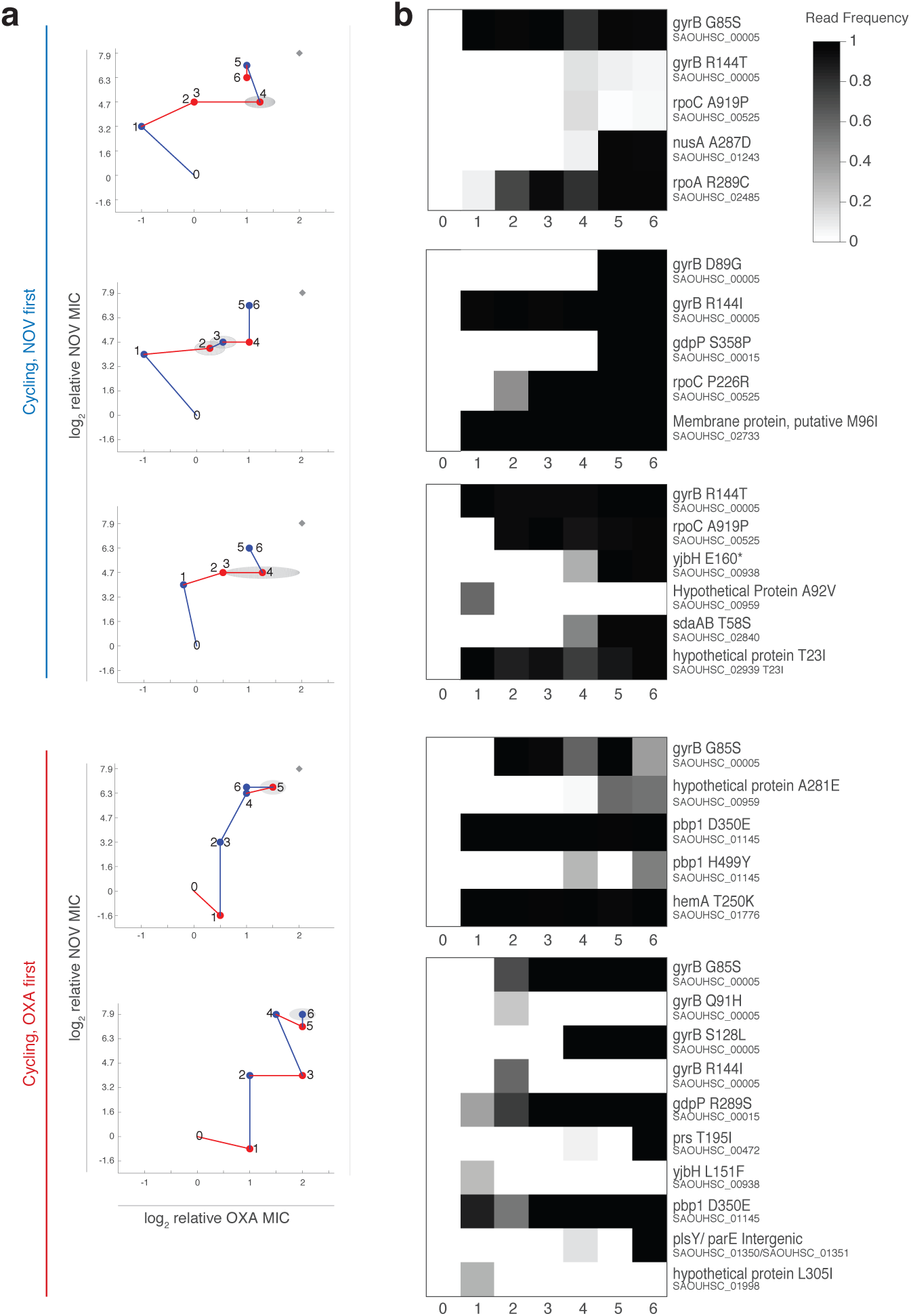
Sequential treatment in OXA and NOV with a strong selective bottleneck suggest that mutations do not interact additively as they accumulate. a) Phenotype changes of 5 replicate populations throughout OXA and NOV selective cycling, as measured by the fold-change of mutant MICs to OXA and NOV. Three replicates were selected first in NOV, and two replicates were first selected in OXA. Numbers indicate steps in the cycling process. Blue lines show a selection step in NOV, while red lines show a selection step in OXA. Gray ellipses illustrate standard deviations from three biological replicates for MIC measurements for both NOV (vertical axis) and OXA (horizontal axis). Gray diamond indicates the median end point of three control populations (cycled between OXA and TSB, or NOV and TSB). b) Genotypic change throughout cycling for each population in (a). Grayscale shading indicates the frequency of reads supporting each mutation, with white as 100%, and black as 0%. Each row represents a single mutation, labeled with the locus tag and amino acid change. Each column is a single step during he cycling process

## Discussion

Our results show that rare positive-cross resistant mutants may circumvent negative cross-resistance. Thus, disease burden and the frequency of resistance mutations should be important factors in the decision to use negative cross-resistance antibiotic therapy in the clinic. Further, even for small population size, antibiotic cycling may not inhibit resistance: the accumulation of negative cross resistance mutations may result in positive cross resistance either additively or due to epistatic interactions, or rare positive cross resistant mutants may arise by chance alone. While the mechanism of negative cross-resistance between OXA and NOV is beyond the scope of this study, the mutations in *gyrB* and *sigB* binding factors suggest that it may be caused by an imbalance between negative supercoiling in DNA and the transcriptional stress response.^34^ However, unraveling the exact mechanisms underlying the diversity of cross-resistance phenotypes will require further study of these mutations in both isolation and combinations.

It would be interesting to see how these results extend to other genetic backgrounds, species, and antibiotics. Ultimately, the successful use of negative cross-resistance cycling to suppress antibiotic resistance will depend on the identification of drugs or drug combinations for which mutual negative cross-resistance is sufficiently frequent even in large populations. Beyond the *de novo* mutations characterized in this study, mobile genetic elements may have their own unique signature of negative and positive cross-resistance. Future designs of sequential antibiotic therapy must take into account the full distribution of possible mutations and their cross-resistance profiles.

## Methods

### Media, strain, and antibiotics

The ancestral strain used for all experiments is *Staphylococcus aureus* RN4220, a phage-free strain derived from NCTC 8325 ^35^. Independent cultures of the ancestor were inoculated with single colonies, grown overnight at 37°C, and stored in 16.7% (v/v) glycerol at -80°C. All experiments were conducted in either trypticase soy agar (TSA, BD 211046) or trypticase soy broth (TSB, BD 211771). Antibiotic solutions were prepared from powder at 10mg/ml in water and stored at 4°C (unless otherwise specified): Oxacillin (Sigma 28221); Novobiocin (Sigma N6160); Ciprofloxacin (Sigma 17850), in 100mM HCl; Doxycycline (Sigma 09891); Gentamycin (Sigma G1264), 50mg/ml; Amikacin (Sigma A1774), 50mg/ml.

### Evolution of first-step mutants

Series of 8-well plate (VWR 267062) were filled with TSA containing increasing concentrations of antibiotic. A serial dilution of the ancestor at 1:10^1.2^ was plated in each well. This ensured that individual colonies can be seen at every antibiotic concentration, and allowed accurate counting of survivors. Plates were incubated at 37°C for 7 days, and imaged using a robotic imaging platform^8^. Automated image segmentation and cell counting in Matlab was reviewed by eye. Frequency of survivors as a function of antibiotic concentration was then calculated. Surviving colonies residing in the MSW were picked using a robotic colony picker (Hudson Robotics, RapidPick) into 96-well microtiter plates (Corning 3628) containing 16.7% (v/v) glycerol in 150uL TSB. Each 96-well plate contains a number of empty wells as negative control and ancestral cells for downstream MIC measurement, such that a combination of four 96-well plates into a 384-well plate will result in 322 wells containing mutants, 16 wells containing the ancestor, and 46 negative controls. 96-well plates were incubated overnight at 37°C, 220 rpm and stored at -80°C.

### High throughput measurement of cross-resistance

Mutant libraries stored in 96-well microtiter plates were replicated with a 96-well floating-pin replicator into a fresh 96-well microtiter plate containing TSB and incubated overnight at 37°C, 220 rpm. Immediately prior to MIC measurement, replicated plates were combined into 384-well microtiter plates (VWR 82030-992) using a 96-channel benchtop pipettor (Rainin 17014207). A total of 16 wildtype cultures from frozen stocks were added to empty wells in the 384-well plate. A robotic liquid handler (Perkin-Elmer SciClone ALH 3000) with a 384-well floating-pin replicator (V&P Scientific) was used to pin 384-well plates onto a TSA plates containing increasing concentrations of antibiotics. Drug serial dilution concentrations for MIC measurement were chosen with 4 steps before the MIC, and 6 steps between the MIC and MPC. TSA plates were incubated at 37°C overnight, before being imaged using a robotic imaging platform. Image segmentation and determination presence or absence of colony was performed in Matlab, followed by manual curation. Cross-resistance scores were calculated as the number of plates that a mutant grew on subtracted by the median number of plates the wildtype grew on. Mutants found to not be resistant to the antibiotic it was selected in were discarded for downstream analysis.

### Construction of directed graphs

A directed, weighted graph was generated using the python library graph-tool^36^, with antibiotics as nodes, and the cross-resistance of each mutant selected in an antibiotic as directed edges originating from that antibiotic.

### Measurement of the impact of population size on negative cross resistance cycling

For selection in OXA followed by plating on NOV, 10^6^ ancestors (low inoculum) were diluted in 5 mL of TSB supplemented with increasing concentrations of OXA. 10^9^ ancestors (high inoculum) were diluted in 100mL of TSB supplemented with the same gradient of OXA. To measure the OXA MIC, 10^3^ ancestors were diluted into a 96-well microtiter plate containing 150uL each of the same gradient used for the low and high inoculum populations. All cultures were incubated overnight at 37°C, 220 rpm. Optical density at 500nm (OD500) was measured with a Perkin-Elmer Victor3, through a 500/20nm emission filter. The OXA concentration at which both high and low inoculum populations were picked was defined as the highest drug concentration in which growth was observed in the 10^6^ population (OD500 above 0.2). Both low and high inoculum cultures were washed twice in TSB and diluted in TSB supplemented with 16.7% glycerol. Aliquots of both populations were stored at -80°C. Population density was measured by thawing aliquots of both populations and performing a serial dilution onto petri dishes containing TSA and incubated overnight at 37°C. Population density was calculated as number of observed colonies divided by the effective volume plated. To measure the impact of inoculum size on population survival, aliquots of high inoculum, low inoculum, and ancestor were plated onto a series of 8-well plates containing increasing concentrations of NOV in TSA, as described in Evolution of first-step mutants. 8-well plates were imaged every day for a total of 7 days on a robotic imaging platform. The time point with maximal difference in population survival was chosen. For the OXA population, this occurred at 7 days. For the NOV population, this occurred at 2 days. Selection in NOV followed by plating in OXA proceeded identically as described, with the exception in that the low inoculum population consists of 10^7^ cells to be consistent with the height of the NOV MSW.

### Whole genome sequencing and analysis

We picked a diverse set of mutants from Fig. 2 (E1 strains) and Fig. 3 (E2 strains). For the E1 set, 5 strains were picked at random from the MSW at each drug concentration where individual colonies were visible. E2 strains were after liquid selection in OXA or NOV. Genomic DNA was isolated using a PureLink 96-well DNA kit (Life Technologies). Library preparation followed ^37^. In short, the DNA was prepared with an Illumina Nextera Kit at reduced volume, with alternate reagents for PCR and DNA purification and custom barcodes for high-density multiplexing. Alignment and mutation calling were performed using breseq ^38^. We wrote a custom wrapper to collate the output of breseq (available at [github]), and performed further downstream analysis in Matlab. Because cycling populations may be a heterogeneous mixture of many clones, we wrote a custom wrapper based on Samtools to count reads supporting all mutations called by breseq in our dataset in all cycling samples, regardless of whether a mutation was originally called in that sample. We were thus able to detect whether mutations called in later cycling steps existed at subclonal frequencies in earlier steps.

### *In vitro* negative cross-resistance cycling

Small populations of ancestor were treated sequentially with OXA and NOV for a total of 6 selection steps. At each step of the cycle, either 10^7^ cells (NOV, one mutation expected) or 10^6^ cells (OXA, one mutation expected) were diluted into 5 mL TSB supplemented with the appropriate antibiotic. The highest drug concentration with OD500 greater than 0.2 was propagated by first performing a sterilizing dilution, and using the dilution step expected to yield either 1 cell (for NOV) or 10 cells (for OXA) to severely bottleneck the population size. Different starting populations sizes are used for OXA and NOV to control for total generation time.

Controls for this experiment consisted of populations that were treated by alternating rounds of NOV and TSB, or OXA and TSB. For the NOV control, 10^7^ cells were propagated in 5 mL NOV gradient or 10^6^ cells were propagated in TSB. For OXA control, 10^6^ cells were propagated in OXA or 10^7^ cells were propagated in NOV.

At each step, aliquots of the propagated population were washed twice in TSB and stored in TSB supplemented with 16.7% glycerol, -80°C. For MIC measurements, frozen aliquots were pinned into 96-well microtiter plates and grown overnight at 37°C, 220 rpm. These populations were then replica pinned into series of 384-well microtiter plates containing increasing concentrations of OXA or NOV and incubated overnight at 37°C, 220 rpm. Optical density at 500nm (OD500) was measured with a Perkin-Elmer Victor3, through a 500/20nm emission filter.

### Statistical testing for enrichment *sigB* binding partners

We performed a two-tailed fisher exact test on a contingency table of all genes in the *Staphylococcus aureus* genome (2892). The margins consist of 58 unique genes found in OXA-resistant mutants and 15 binding partners of *sigB*^28^. Columns indicate whether genes are *sigB* binding partners, and rows indicate whether genes were mutated in our dataset. Two-tailed fisher exact test gave p-value of 1.69⋅10^-4^.

### Statistical testing for association of mutation with cross-resistance phenotype

To limit the number of tests, we eliminated all events that occurred fewer than 3 times in all phenotypes. We then performed two-tailed fisher exact tests to detect associations between each specific mutation and cross-resistance phenotype.

## References

1. Centers for Disease Control and Prevention. Antibiotic Resistance Threats in the United States, 2013. 1–114 (2013).

2. World Health Organization. Antimicrobial resistance global report on surveillance : 2014 summary. (2014).

3. Baym, M., Stone, L. K. & Kishony, R. Multidrug evolutionary strategies to reverse antibiotic resistance. Science 351, aad3292 (2016).

4. Szybalski, W. & Bryson, V. Genetic studies on microbial cross resistance to toxic agents. I. Cross resistance of Escherichia coli to fifteen antibiotics. J Bacteriol 64, 489–499 (1952).

5. Imamovic, L. & Sommer, M. O. A. Use of collateral sensitivity networks to design drug cycling protocols that avoid resistance development. Sci Transl Med 5, 204ra132 (2013).

6. Lázár, V. et al. Bacterial evolution of antibiotic hypersensitivity. Molecular Systems Biology 9, 700–700 (2013).

7. Kim, S., Lieberman, T. D. & Kishony, R. Alternating antibiotic treatments constrain evolutionary paths to multidrug resistance. Proc Natl Acad Sci USA 111, 14494–14499 (2014).

8. Stone, L. K., Baym, M., Lieberman, T. D. & Chait, R. Compounds that select against the tetracycline-resistance efflux pump. Nature Chemical … (2016).

9. Lázár, V. et al. Genome-wide analysis captures the determinants of the antibiotic cross-resistance interaction network. Nat Commun 5, 4352 (2014).

10. Lieberman, T. D. et al. Parallel bacterial evolution within multiple patients identifies candidate pathogenicity genes. Nat Genet 43, 1275–1280 (2011).

11. Musher, D. M. et al. Emergence of Macrolide Resistance during Treatment of Pneumococcal Pneumonia. New Engl J Med 346, 630–631 (2002).

12. Lipsitch, M. & Levin, B. R. The population dynamics of antimicrobial chemotherapy. Antimicrobial agents and chemotherapy 41, 363–373 (1997).

13. Stone, N. R. H. et al. Breakthrough bacteraemia due to tigecycline-resistant Escherichia coli with New Delhi metallo-lactamase (NDM)-1 successfully treated with colistin in a patient with calciphylaxis. Journal of Antimicrobial Chemotherapy 66, 2677–2678 (2011).

14. Oz, T. et al. Strength of selection pressure is an important parameter contributing to the complexity of antibiotic resistance evolution. Mol Biol Evol 31, 2387–2401 (2014).

15. Dragosits, M., Mozhayskiy, V., Quinones-Soto, S., Park, J. & Tagkopoulos, I. Evolutionary potential, cross-stress behavior and the genetic basis of acquired stress resistance in Escherichia coli. Molecular Systems Biology 9, 643–643 (2013).

16. Perron, G. G., Gonzalez, A. & Buckling, A. Source-sink dynamics shape the evolution of antibiotic resistance and its pleiotropic fitness cost. Proc. Biol. Sci. 274, 2351–2356 (2007).

17. Gressel, J. & Segel, L. A. Negative cross-resistance; a possible key to atrazine resistance management: a call for whole plant data. Z Naturforsch. 45c, 470–473 (1990).

18. Gadamski, G., Ciarka, D., Gressel, J. & Gawronski, S. W. Negative cross-resistance in triazine-resistant biotypes of Echinochloa crus-galliand Conyza canadensis. Weed Science 48, 176–180 (2000).

19. de Evgrafov, M. R., Gumpert, H., Munck, C., Thomsen, T. T. & Sommer, M. O. A. Collateral resistance and sensitivity modulate evolution of high-level resistance to drug combination treatment in Staphylococcus aureus. Mol Biol Evol 32, msv006–1185 (2015).

20. Pál, C., Papp, B. & Lázár, V. Collateral sensitivity of antibiotic-resistant microbes. Trends in Microbiology 23, 1–7 (2015).

21. Ba, X. et al. Novel mutations in penicillin-binding protein genes in clinical Staphylococcus aureus isolates that are methicillin resistant on susceptibility testing, but lack the mec gene. Journal of Antimicrobial Chemotherapy 69, 594–597 (2014).

22. Howden, B. P., Davies, J. K., Johnson, P. D. R., Stinear, T. P. & Grayson, M. L. Reduced Vancomycin Susceptibility in Staphylococcus aureus, Including Vancomycin-Intermediate and Heterogeneous Vancomycin-Intermediate Strains: Resistance Mechanisms, Laboratory Detection, and Clinical Implications. Clin. Microbiol. Rev. 23, 99–139 (2010).

23. Mwangi, M. M. et al. Tracking the in vivo evolution of multidrug resistance in Staphylococcus aureus by whole-genome sequencing. Proc Natl Acad Sci USA 104, 9451–9456 (2007).

24. Tsiodras, S. et al. Linezolid resistance in a clinical isolate of Staphylococcus aureus. The Lancet 358, 207–208 (2001).

25. Gao, W. et al. Two Novel Point Mutations in Clinical Staphylococcus aureus Reduce Linezolid Susceptibility and Switch on the Stringent Response to Promote Persistent Infection. PLoS Pathog 6, e1000944 (2010).

26. Drlica, K. & Zhao, X. Mutant Selection Window Hypothesis Updated. Clinical Infectious Diseases 44, 681–688 (2007).

27. Stieger, M., Angehrn, P., Wohlgensinger, B. & Gmünder, H. GyrB mutations in Staphylococcus aureus strains resistant to cyclothialidine, coumermycin, and novobiocin. Antimicrobial agents and chemotherapy 40, 1060–1062 (1996).

28. Cherkasov, A. et al. Mapping the protein interaction network in methicillin-resistant Staphylococcus aureus. J. Proteome Res. 10, 1139–1150 (2011).

29. Corrigan, R. M., Abbott, J. C., Burhenne, H., Kaever, V. & Gründling, A. c-di-AMP is a new second messenger in Staphylococcus aureus with a role in controlling cell size and envelope stress. PLoS Pathog 7, e1002217 (2011).

30. Griffiths, J. M. & O'Neill, A. J. Loss of function of the gdpP protein leads to joint β-lactam/glycopeptide tolerance in Staphylococcus aureus. Antimicrobial agents and chemotherapy 56, 579–581 (2012).

31. Dengler, V. et al. Mutation in the C-di-AMP cyclase dacA affects fitness and resistance of methicillin resistant Staphylococcus aureus. PLoS ONE 8, e73512 (2013).

32. Komatsuzawa, H. et al. Cloning and characterization of the fmt gene which affects the methicillin resistance level and autolysis in the presence of triton X-100 in methicillin-resistant Staphylococcus aureus. Antimicrobial agents and chemotherapy 41, 2355–2361 (1997).

33. Miyazaki, E., Chen, J. M., Ko, C. & Bishai, W. R. The Staphylococcus aureus rsbW (orf159) gene encodes an anti-sigma factor of SigB. J Bacteriol 181, 2846–2851 (1999).

34. Rovinskiy, N., Agbleke, A. A., Chesnokova, O., Pang, Z. & Higgins, N. P. Rates of gyrase supercoiling and transcription elongation control supercoil density in a bacterial chromosome. PLOS Genet 8, e1002845 (2012).

35. Nair, D. et al. Whole-genome sequencing of Staphylococcus aureus strain RN4220, a key laboratory strain used in virulence research, identifies mutations that affect not only virulence factors but also the fitness of the strain. J Bacteriol 193, 2332–2335 (2011).

36. Peixoto, T. P. The graph-tool python library. figshare (2014), doi:10.6084/m9.figshare.1164194 (2014). doi:10.6084/m9.figshare.1164194

37. Baym, M. et al. Inexpensive multiplexed library preparation for megabase-sized genomes. PLoS ONE 10, e0128036 (2015).

38. Deatherage, D. E. & Barrick, J. E. Identification of mutations in laboratory-evolved microbes from next-generation sequencing data using breseq. Methods Mol. Biol. 1151, 165–188 (2014).

39. Fujimoto-Nakamura, M., Ito, H., Oyamada, Y., Nishino, T. & Yamagishi, J.-I. Accumulation of mutations in both gyrB and parE genes is associated with high-level resistance to novobiocin in Staphylococcus aureus. Antimicrobial agents and chemotherapy 49, 3810–3815 (2005).

